# Anti-Nucleolin Aptamer iSN04 Inhibits Alveolar Rhabdomyosarcoma Cell Growth by Inducing Myogenic Differentiation

**DOI:** 10.64898/2026.10.24.761494

**Authors:** Yoshihiro Saitoh, Tomohide Takaya

**Affiliations:** Department of Agriculture, Graduate School of Medicine, Science and Technology, Shinshu University, 8304 Minami-minowa, Kami-ina, Nagano 399-4598, Japan; Department of Agricultural and Life Sciences, Faculty of Agriculture, Shinshu University, 8304 Minami-minowa, Kami-ina, Nagano 399-4598, Japan

**Keywords:** alveolar rhabdomyosarcoma, aptamer, myogenetic oligodeoxynucleotide, nucleolin, vincristine

## Abstract

Rhabdomyosarcoma (RMS) is a muscle-derived soft tissue tumor that retains myogenic potential. Novel drugs for RMS are strongly needed because the standard chemotherapeutic agents for RMS, including vincristine (VCR), have marginally improved the survival rate of high-risk patients for several decades. An anti-nucleolin aptamer, iSN04, was originally identified to promote myogenic differentiation of myoblasts, and was subsequently reported to suppress the growth of embryonal RMS cells by potentiating their differentiation and arresting proliferation. The present study investigated the effects of iSN04 on two alveolar RMS cell lines, NRS-1 and RH30. iSN04 modulated gene expression levels such as myogenic transcription factors (MyoD and myogenin), inhibitor of differentiation proteins (ID1 and ID3), and the tumor suppressor CASZ1. iSN04 treatment induced NRS-1 cells to differentiate into contracting myotube-like cells, and severely impaired the ability of RH30 cells to form and grow tumorspheres in three-dimensional culture. Co-treatment with iSN04 and a low dose of VCR cooperatively suppressed the proliferation of RH30 cells compared with single-agent treatments. These data demonstrate that iSN04 can be a beneficial agent for RMS and hold potential for use in combination therapy with current standard drugs.

## 1. Introduction

Rhabdomyosarcoma (RMS) is a striated muscle-derived soft-tissue sarcoma, representing the most common soft-tissue tumor in children. The two major subtypes are embryonal RMS (ERMS, accounting for 60-70% of cases) and alveolar RMS (ARMS, 20-30%) [Hebron, 2026]. ERMS typically arises from causative mutations in genes such as *TP53* within myogenic lineages, such as skeletal muscle stem cells (satellite cells) and myogenic precursor cells (myoblasts) [Rubin 2011]. In contrast, most ARMS cases are associated with chromosomal translocations expressing the fusion oncogene *PAX3::FOXO1* [Barr 1993] or *PAX7::FOXO1* [Davis 1994]. Standard multidisciplinary treatment for RMS comprises chemotherapy, surgical resection, and radiation therapy. Multi-drug chemotherapy using vincristine (VCR), actinomycin D (ActD), and cyclophosphamide/ifosfamide has cured the patients with localized RMS [Miwa 2022]. However, the five-year survival rate of highrisk patients with metastatic RMS has remained at 20-30% for several decades [Haduong 2022]. VCR, which interferes with the polymerization of tubulin and its incorporation into microtubules, has been used as an anti-cancer drug. However, VCR treatment is limited by its side effects, such as neuropathy [Madsen 2019]. High-dose chemotherapy for RMS patients has been reported to increase the incidence of treatment-related adverse events [Carli 2004]. Thus, novel drugs need to be developed to improve the therapeutic strategy for RMS. Differentiation therapy has been studied based on the findings that RMS cells resemble poorly differentiated muscle cells. Trametinib, a MEK inhibitor, was reported to induce myogenic differentiation of RMS cells, suppress tumor growth, and prolong overall survival in animal models [Yohe 2018, Hebron 2025].

We have previously reported that a lactic acid bacterium genome-derived 18-mer oligodeoxynucleotide, iSN04 (5’-AGA TTA GGG TGA GGG TGA-3’), serves as an anti-nucleolin aptamer and robustly induces myoblast differentiation and myotube formation [Shinji 2021]. Nucleolin, the target of iSN04, is a multifunctional phosphoprotein that is ubiquitously expressed and localizes in the nucleolus, nucleoplasm, cytoplasm, and plasma membrane. Nucleolin interacts with various binding partners and is involved in numerous cellular processes, such as ribosome biogenesis, nucleic acid metabolism, apoptosis, cell proliferation, and differentiation [Jia 2017]. Nucleolin has been reported to be overexpressed in tumor cells to promote tumorigenesis [Prakash 2025], including that of RMS [Willmer 2021]. It is widely accepted that nucleolin can serve as a therapeutic target or diagnostic marker for tumors [Berger 2015]. We previously demonstrated that iSN04 effectively inhibits the growth of three ERMS cell lines, ICH-ERMS1, KYM1, and RD, by the partial promotion of myogenic differentiation [Nohira 2022]. These findings indicated that iSN04 could be a beneficial agent for the differentiation therapy for RMS. However, the anti-RMS effect of iSN04 on ARMS cells has not yet been elucidated.

This study investigated whether iSN04 induces myogenic differentiation and suppresses the proliferation of two ARMS cell lines, NRS-1 and RH30. Furthermore, the combination effect of iSN04 and VCR was evaluated to assess the feasibility of iSN04 in clinical applications.

## 2. Materials and Methods

### 2.1. Chemicals

iSN04, in which all phosphodiester bonds were phosphorothioated to enhance nuclease resistance in cell culture, was synthesized and HPLC-purified (GeneDesign, Osaka, Japan). iSN04 and VCR sulfate salt (Merck, Darmstadt, Germany) were dissolved in endotoxin-free water. An equal volume of endotoxin-free water was used as a negative control.

### 2.2. Cell Culture

Two *PAX3-FOXO1*-positive cell lines, NRS-1 and RH30, were used. NRS-1, derived from the left forearm ARMS of a 7-year-old female [Ogose 1995], was provided by the RIKEN BRC Cell Bank (RCB1188; BioResource Research Center, RIKEN, Tsukuba, Japan); RH30, derived from the muscle ARMS of a 17-year-old male [Hinson 2013], was provided by the ATCC (CRL-2061; American Type Culture Collection, Manassas, VA, USA). Both NRS-1 and RH30 cells were maintained in culture medium consisting of RPMI 1640 (Nacalai, Osaka, Japan), 10% fetal bovine serum (FBS) (HyClone; GE Healthcare, Chicago, IL, USA), and a mixture of 100 units/ml penicillin and 100 μg/ml streptomycin (P/S) (Nacalai). All cells were cultured at 37°C with 5% CO_2_ throughout the experiments.

### 2.2. Intracellular Immunocytochemistry

NRS-1 and RH30 cells were seeded in 30-mm dishes (2.0×10^5^ cells/dish), fixed with 2% paraformaldehyde, permeabilized with 0.2% Triton X-100, immunoreacted with 1.0 μg/mL rabbit polyclonal anti-nucleolin antibody (ab22758; Abcam, Cambridge, UK) or 0.5 μg/mL mouse monoclonal anti-pan-myosin heavy chain (MHC) antibody (MF20; R&D Systems, Minneapolis, MN, USA), and then immunostained with 0.1 μg/ml of Alexa Fluor 488-conjugated donkey polyclonal anti-rabbit or anti-mouse IgG antibody (Jackson ImmunoResearch, West Grove, PA, USA). Cell nuclei were stained with 4’,6-diamidino-2-phenylindole (DAPI) (Nacalai). Fluorescence images were captured using an EVOS FL Auto microscope (AMAFD1000; Thermo Fisher Scientific, Waltham, MA, USA).

### 2.3. Induction of Myogenic Differentiation

NRS-1 cells were seeded in 30-mm dishes (2.0×10^5^ cells/dish) and were induced to undergo myogenic differentiation in differentiation medium consisting of DMEM (Nacalai), 2% horse serum (HyClone; Cytiva, Marlborough, MA, USA), and P/S. After 48 h of seeding, the cells were treated with 10 μM iSN04. After 7 days of treatment, phasecontrast videos were captured using a CKX53 microscope (Olympus, Tokyo, Japan) equipped with a Moticam 1080 digital camera system (Shimadzu RIKA Corporation, Tokyo, Japan).

### 2.4. Three-Dimensional (3D) Culture of Tumorsphere

RH30 cells were dissociated and suspended in culture medium. Droplets of cell suspension (3000 cells/30 μl) were placed on 24-well floating culture plates (Sumitomo Bakelite, Tokyo, Japan). The plates were then inverted for hanging-drop culture. After 3 days, the plates were inverted again, and 300 μl/well of culture medium containing 10 μM iSN04 was added to the RH30 tumorspheres (designated as day 0). The spheres were maintained for 4 days without medium replacement [Nohira 2022, Shinji 2020]. Bright-field images were captured using an EVOS FL Auto microscope.

### 2.4. Quantitative Real-Time RT-PCR (qPCR)

NRS-1 cells were seeded in 60-mm dishes (1.0 × 10^6^ cells/dish) and treated with 10 μM iSN04 for 48 h. RH30 cells were seeded in 30-mm dishes (1.0 × 10^5^ cells/dish) and treated with 10 μM iSN04 and 0.5 nM VCR for 96 h. Total RNA was isolated from the cells using a NucleoSpin RNA Plus kit (Macherey-Nagel, Düren, Germany) and reverse-transcribed using ReverTra Ace qPCR RT Master Mix (TOYOBO, Osaka, Japan). qPCR was performed using GoTaq qPCR Master Mix (Promega, Madison, WI, USA) on a Thermal Cycler Dice Real Time System III (TaKaRa Bio, Kusatsu, Japan). The relative expression level of each transcript was normalized to that of the tyrosine 3-monooxygenase/tryptophan 5-monooxygenase activation protein zeta (*YWHAZ*) gene. Results are expressed as fold changes. Primer sequences are listed in Table 1.

**Table 1.**
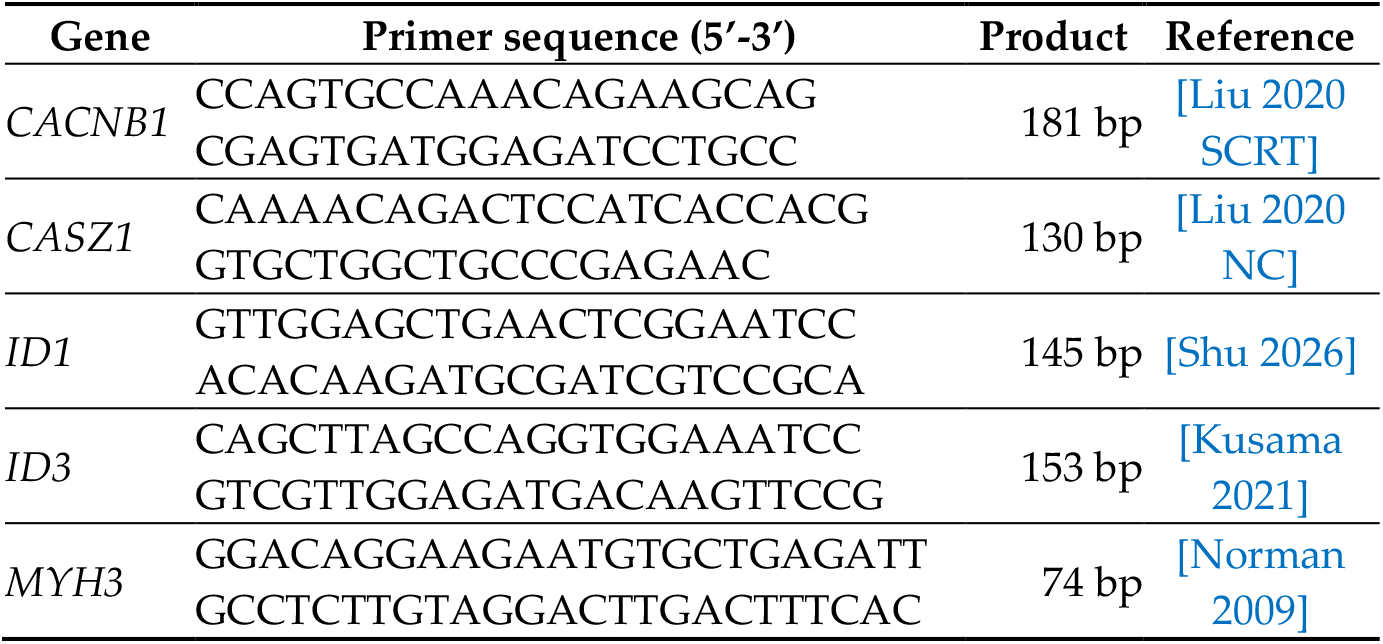
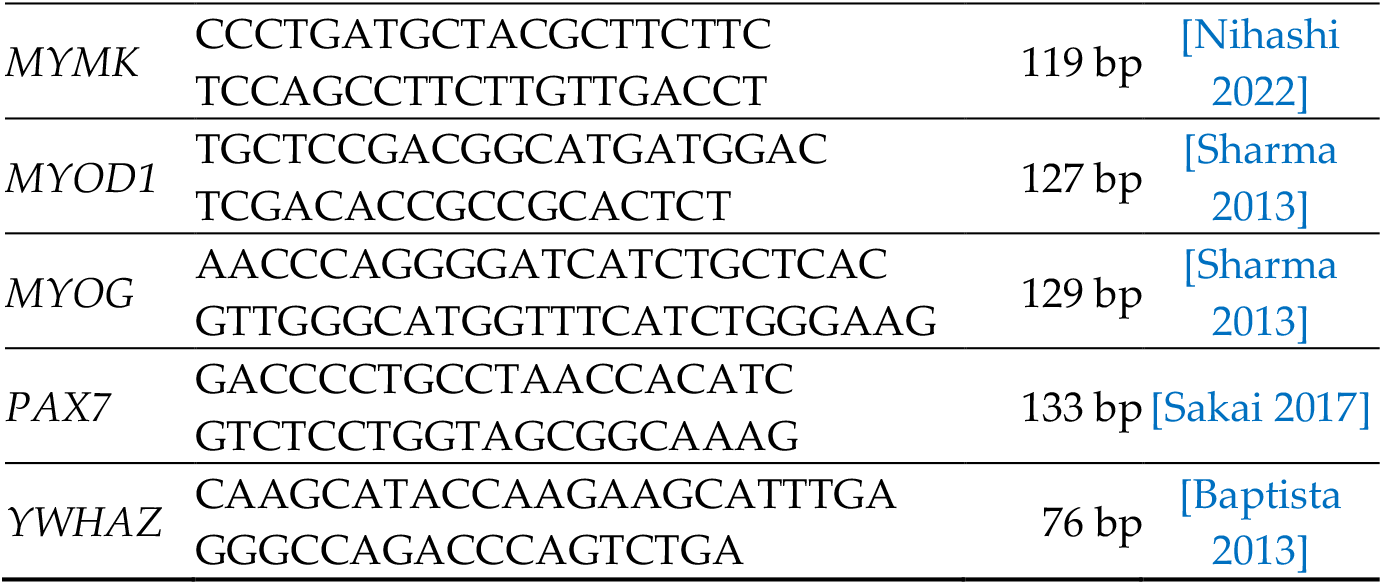
Primer sequences for qPCR.

### 2.5. Cell Counting

NRS-1 cells were seeded in 12-well plates (5.0×10^4^ cells/well) and treated with 10 or 30 μM iSN04 for 96 h. RH30 cells were seeded in 12-well plates (3.0×10^4^ cells/well) and treated with 10 μM iSN04 and 0.5 nM VCR for 96 h. Bright-field images were captured using an EVOS FL Auto microscope every 24 h. When the control group reached confluence, the cells were completely dissociated with 0.25% trypsin and counted using a hemocytometer [Nohira 2022, Shinji 2020].

### 2.6. Statistical Analysis

Results are presented as the mean ± standard error of the mean. Statistical comparisons between two groups were performed using an unpaired two-tailed Student’s *t*-test, and those among multiple groups were performed using the Tukey-Kramer test following a one-way analysis of variance. Statistical significance was set at *p* < 0.05.

## 3. Results

### 3.1. Nucleolin Expression and Myotube Formation of NRS-1 Cells

The NRS-1 cell line has been reported to exhibit spontaneous myogenic differentiation into multinucleated myotube-like cells with myoglobin expression in vitro [Ogose 1995]. Although NRS-1 cells have been used in some studies as an ARMS cell line [Yoshitaka 2013, Jaaks 2016, Kato 2021, Ishiguro 2024], their myogenic potential has not been elucidated in detail.

Intracellular immunostaining revealed that nucleolin localized to the nucleoplasm and intensively to the nucleoli of NRS-1 cells (Figure 1A), resembling that observed in myoblasts [Shinji 2021] and ERMS cells [Nohira 2022]. Immunostaining for MHC, a terminal differentiation marker of skeletal muscle, clearly visualized multinucleated myotube-like morphology with sarcomere-like structures in NRS-1 cells (Figure 1B). These results suggest that antagonizing nucleolin with iSN04 would promote the myogenic differentiation of NRS-1 cells, as previously reported in myoblasts [Shinji 2021] and ERMS cells [Nohira 2022].

**Figure 1.**
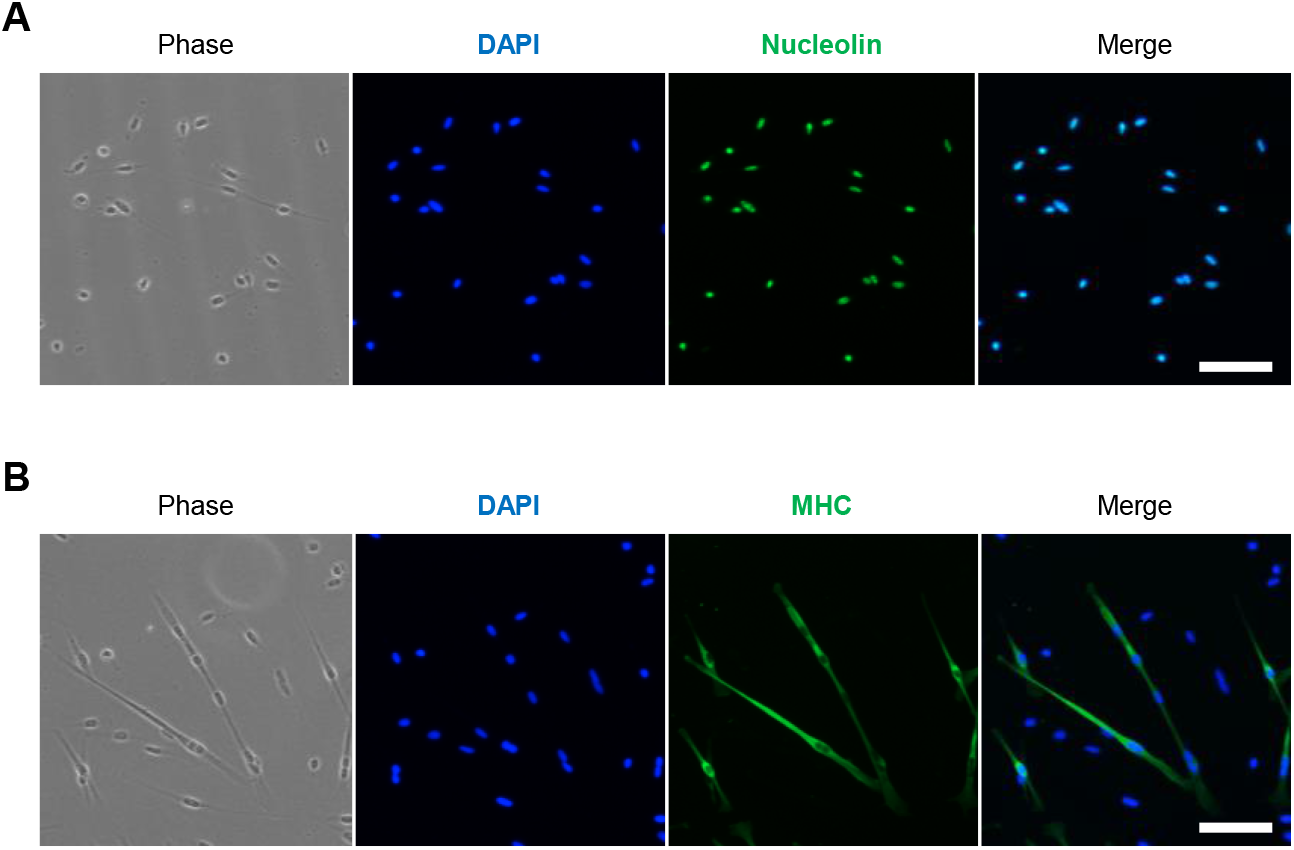
Representative fluorescence images of immunostaining for nucleolin (**A**) and MHC (**B**) in NRS-1 cells. Scale bars, 100 μm.

NRS-1 cells were induced to undergo myogenic differentiation in differentiation medium with 10 μM iSN04. After 7 days of treatment, spontaneously contracting myotube-like cells were observed in the iSN04-treated group but not in the control group (Supplementary Video S1). This demonstrates that iSN04 induces myogenic differentiation, myotube formation, and/or myotube maturation of NRS-1 cells by inhibiting nucleolin.

### 3.2. iSN04 Induces Myogenic Differentiation of NRS-1 Cells

The MHC-positive, multinucleated, and contracting myotube-like cells observed in the iSN04-treated NRS-1 cells suggest that myogenic potential is relatively well preserved in NRS-1 cells. Therefore, we investigated how iSN04 restores the myogenic potential of NRS-1 cells.

qPCR revealed that iSN04 modulates myogenic gene expression in NRS-1 cells (Figure 2A). iSN04 treatment did not alter the expression levels of Pax7 and embryonic MHC (*MYH3*). Although there was no significant difference, iSN04 consistently upregulated the mRNA levels of myogenic genes such as MyoD (*MYOD1*), myogenin (*MYOG*), myomaker (*MYMK*), and voltage-gated calcium channel β1 subunit (Ca_v_β^1^) (*CACNB1*). MyoD and myogenin are myogenic regulatory factors (MRFs) serving as transcription factors to promote muscle gene expression. Myomaker is an MRF-driven muscle-specific transmembrane protein required for myoblast fusion to form multinucleated myotubes [Zhang 2020]. Ca_v_β^1^, a subunit of the dominant calcium channel in muscle cells, plays an essential role in excitation-contraction coupling [Shishmarev 2020] and enters the nucleus to regulate the myogenic gene expression program [Taylor 2014]. The iSN04-induced expression of MyoD, myogenin, and myomaker in NRS-1 cells is consistent with that verified in myoblasts [Shinji 2021, Umezawa 2024] and ERMS cells [Nohira 2022]. We have also confirmed that iSN04 significantly induces *CACNB1* expression in an ERMS cell line, RD (unpublished data). These results and findings demonstrate that iSN04 drives NRS-1 cells to form contracting myotube-like cells by activating the myogenic gene expression program. In addition, iSN04 significantly reduced the mRNA levels of ID1 and ID3 in NRS-1 cells. ID1 and ID3 inhibit MRF activity by competitive binding to E-proteins during myogenesis [Yokoyama 2011]. The reduction of ID1 and ID3 expression by iSN04 is considered to enhance the myogenic differentiation of NRS-1 cells. iSN04 also significantly upregulated CASZ1 expression in NRS-1 cells. CASZ1 is a zinc finger transcription factor that upregulates MyoD signaling. Importantly, loss of CASZ1 activity impairs myogenic differentiation and promotes tumorigenesis of ERMS, whereas restoration of CASZ1 reverses these ERMS phenotypes [Liu 2020 NC]. This strongly suggests that the induction of CASZ1 by iSN04 is part of the mechanism driving the myogenic differentiation of NRS-1 cells.

**Figure 2.**
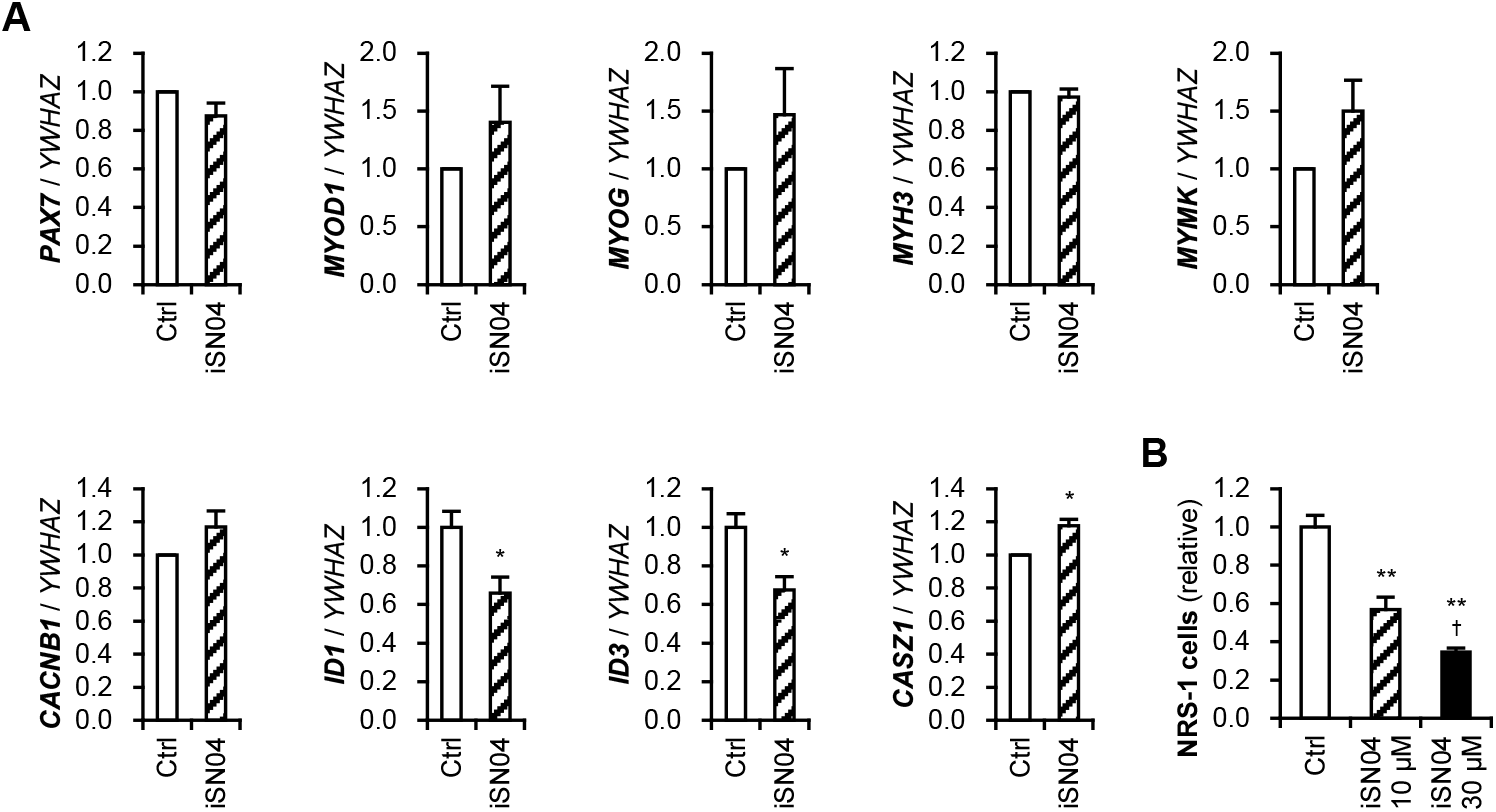
(**A**) qPCR results of NRS-1 cells treated with 10 μM iSN04 for 48 h. * *p* < 0.05 vs. control (Student’s *t*-test). *n* = 3. (**B**) Number of NRS-1 cells treated with 10 or 30 μM iSN04 for 96 h. ** *p* < 0.01 vs. control, ^†^ *p* < 0.05 vs. 10 μM iSN04 (Tukey-Kramer test). *n* = 4.

During myogenesis, myogenic progenitor cells permanently exit the cell cycle and enter terminal differentiation [Singh 2013, Wu 2024]. Therefore, the induction of myogenic differentiation generally suppresses cell proliferation, which underpins the strategy of differentiation therapy for RMS [Yohe 2018, Hebron 2025]. iSN04 treatment significantly reduced the number of NRS-1 cells in a dose-dependent manner (Figure 2B), indicating that iSN04-induced differentiation results in the suppression of NRS-1 cell proliferation.

### 3.3. Nucleolin Expression and Myotube Formation of RH30 Cells

The RH30 cell line is one of the major ARMS cell lines [Hinson 2013] and expresses nucleolin on its surface [Dzhumashev 2024]; however, the intracellular localization of nucleolin and the effect of nucleolin inhibition have not been investigated. Intracellular immunostaining revealed that nucleolin localized to the nucleoplasm and intensively to the nucleoli of RH30 cells, similar to NRS-1 cells (Figure 3A). However, MHC was not detected in RH30 cells at all (Figure 3B), indicating the lower myogenic potential of RH30 cells compared with NRS-1 cells.

**Figure 3.**
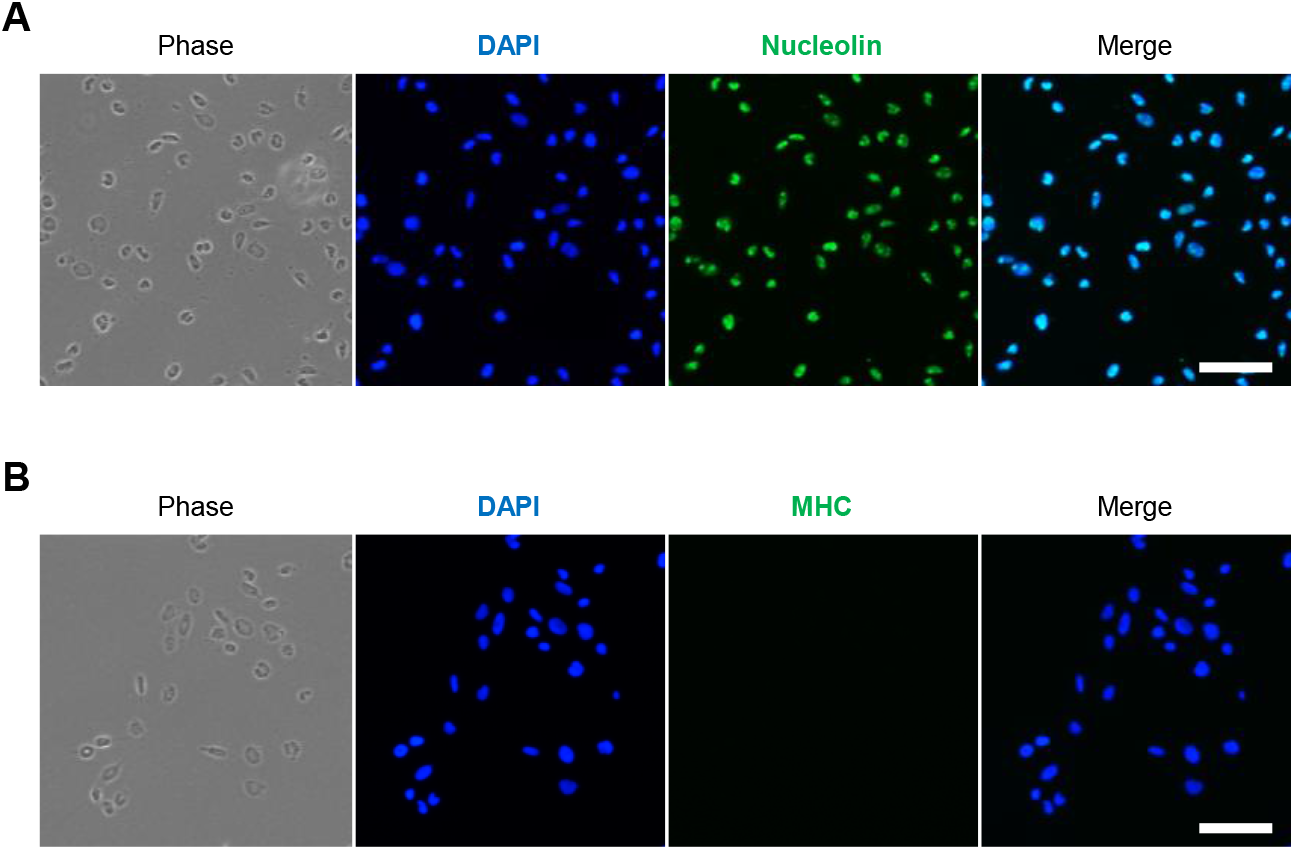
Representative fluorescence images of immunostaining for nucleolin (**A**) and MHC (**B**) in RH30 cells. Scale bars, 100 μm.

### 3.4. iSN04 Disturbs Tumorsphere Formation of RH30 Cells

The effect of iSN04 on RH30 cells was investigated using 3D-cultured tumorspheres to mimic in vivo tumorigenesis [He 2017]. The RH30 tumorspheres formed by hanging-drop culture were subjected to floating culture with iSN04 (Figure 4). In the control group, RH30 tumorspheres grew into globes over 0.5 mm in diameter with adherent outgrowth. However, in the presence of iSN04, RH30 tumorspheres were unable to grow or maintain cell-cell adhesion, eventually collapsing into single cells. These results demonstrate that iSN04 is potentially beneficial for suppressing the growth of ARMS tumors, particularly in their initial stages.

**Figure 4.**
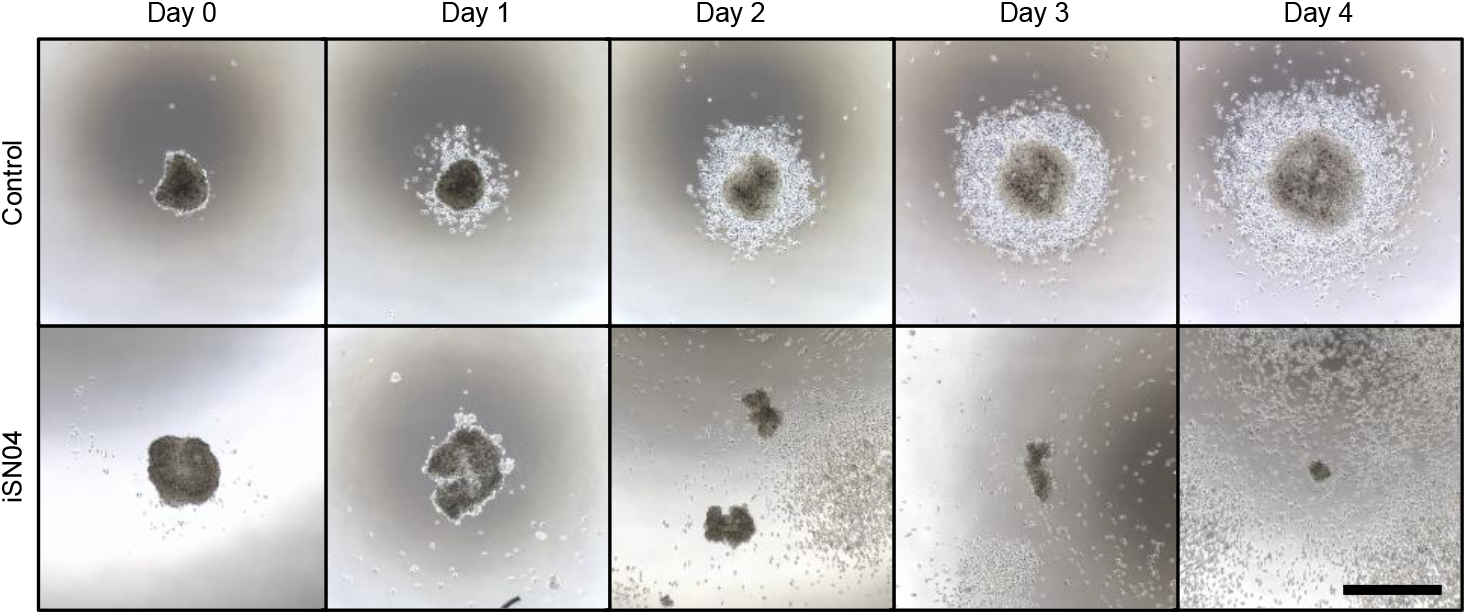
Representative bright-field images of RH30 tumorspheres treated with 10 μM iSN04. Scale bar, 500 μm.

### 3.5 iSN04 But Not VCR Induce Myogenic Gene Expression in RH30 Cells

We investigated whether iSN04 induces myogenic differentiation of RH30 cells. To evaluate the potential of iSN04 in clinical settings, it was co-administered with VCR, one of the standard chemotherapeutic agents for RMS [Miwa 2020]. VCR alone has generally been administered to RH30 cells at a concentration of ~50 nM [Yoon 2020]. However, in combination experiments using RH30 cells, VCR is used at a lower concentration of ~0.5 nM [Urla 2022]. In this study, RH30 cells were treated with 10 μM iSN04 and/or 0.5 nM VCR, and then subjected to qPCR (Figure 5).

**Figure 5.**
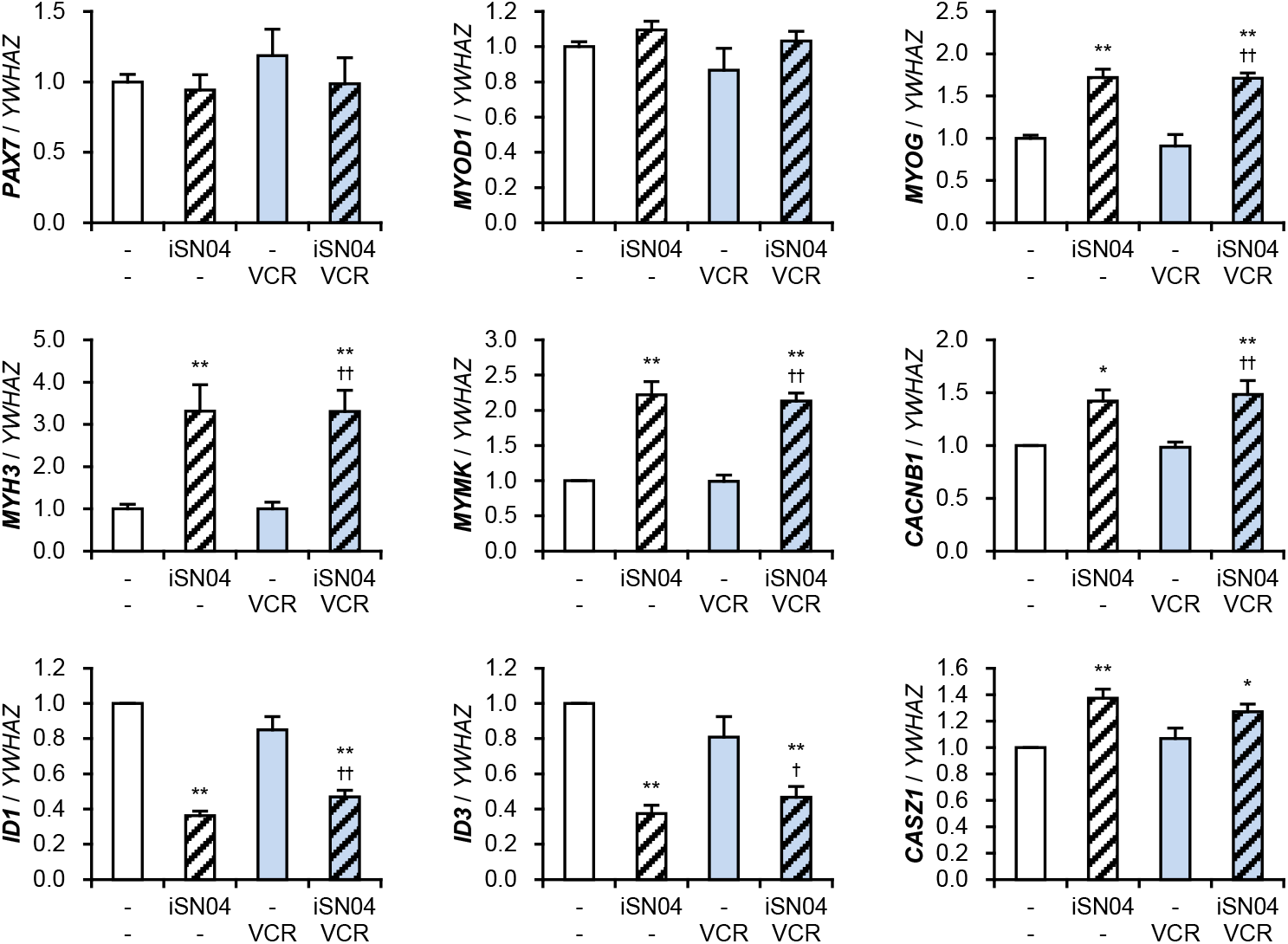
qPCR results of RH30 cells treated with 10 μM iSN04 and 0.5 nM VCR for 96 h. * *p* < 0.05, ** *p* < 0.01 vs. control; ^†^ *p* < 0.05, ^††^ *p* < 0.01 vs. VCR (Tukey-Kramer test). *n* = 4.

VCR alone did not alter the expression levels of any tested genes, demonstrating that

0.5 nM VCR is unable to induce myogenic differentiation of RH30 cells. iSN04 did not alter the mRNA levels of Pax7 and MyoD, but significantly induced the expression of myogenin, embryonic MHC, myomaker, Ca_v_β^1^, and CASZ1. iSN04 also significantly suppressed the expression of ID1 and ID3. These iSN04-dependent changes in gene expression in RH30 cells correspond to those observed in NRS-1 cells (Figure 2A). VCR did not affect the iSN04-dependent alterations in mRNA levels in RH30 cells; there were no significant differences between the iSN04 group and the iSN04 + VCR combination group. These results indicate that iSN04 can induce the differentiation of RH30 cells, which exhibit lower myogenic potential compared with NRS-1 cells. Because VCR does not impair the myogenic effect of iSN04, their drug actions are expected to be independent and beneficial in combination therapy.

### 3.6. iSN04 and VCR Cooperatively Suppress Proliferation of RH30 Cells

The combination effect of iSN04 and VCR on the proliferation of RH30 cells was investigated by sequential observation and cell counting at the end point (Figure 6). The number of the RH30 cells treated with 10 μM iSN04 or 0.5 nM VCR alone was significantly reduced, demonstrating that these single treatments suppress the proliferation of RH30 cells. Importantly, the cell number in the iSN04 + VCR combination group was significantly reduced compared with that of the VCR group. This cooperative effect of iSN04 and VCR on the proliferation of RH30 cells demonstrates that iSN04 is a potential drug candidate to be added to the standard chemotherapy with VCR.

**Figure 6.**
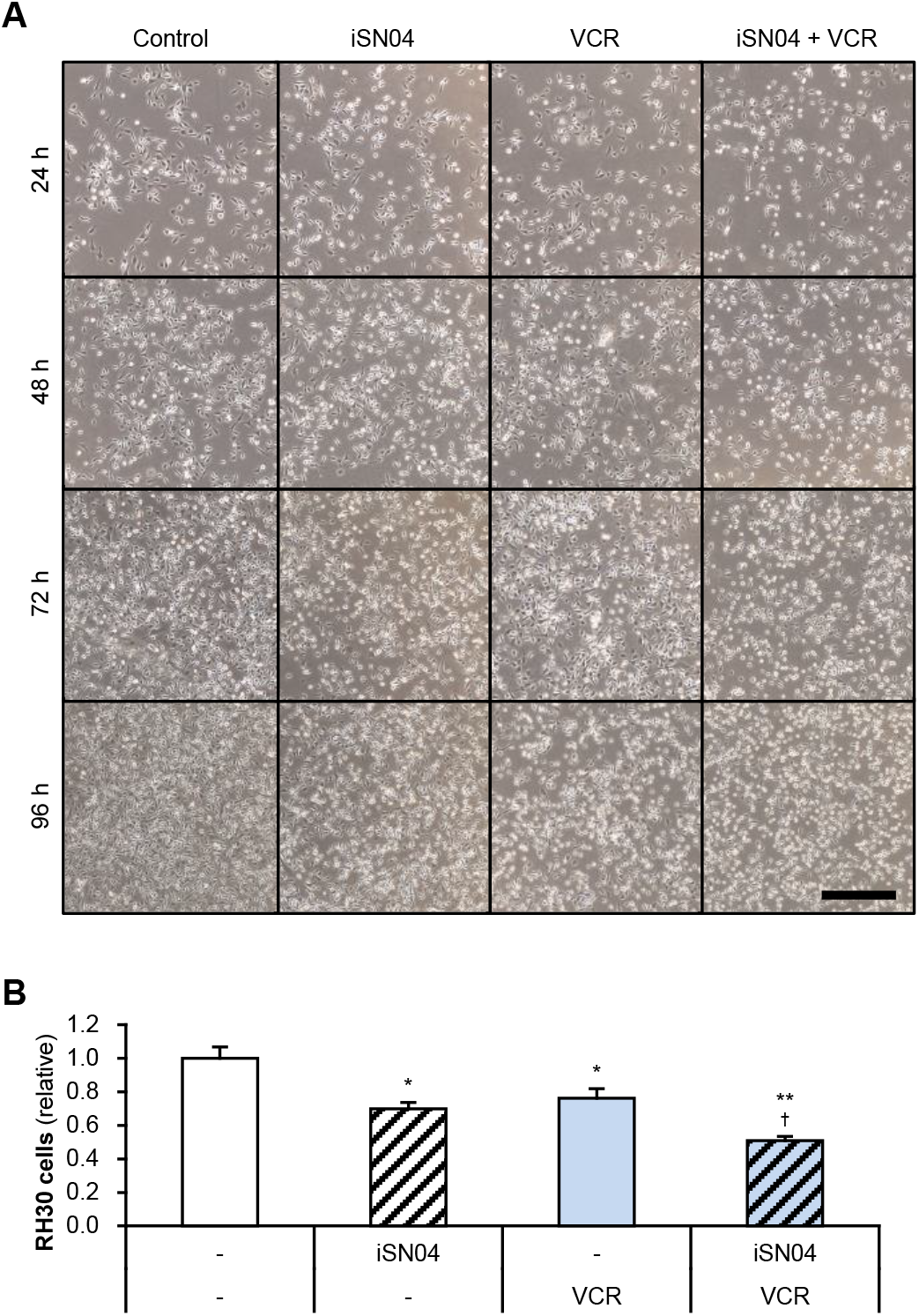
(**A**) Representative phase-contrast images of RH30 cells treated with 10 μM iSN04 and 0.5 nM VCR. Scale bar, 500 μm. (**B**) Number of RH30 cells treated with 10 μM iSN04 and 0.5 nM VCR for 96 h. * *p* < 0.05, ** *p* < 0.01 vs. control; ^†^ *p* < 0.05 vs. VCR (Tukey-Kramer test). *n* = 3.

## 4. Discussion

The present study showed that the anti-nucleolin aptamer, iSN04, induces myogenic differentiation and suppresses the proliferation of two ARMS cell lines, NRS-1 and RH30, in addition to three ERMS cell lines previously reported [Nohira 2022]. These findings demonstrate that iSN04 is a promising candidate nucleic acid drug for differentiation therapy for RMS. Nucleolin is known to contribute to cell proliferation via RNA metabolism and chromatin remodeling [Jia 2017]. In our previous study, antagonizing nucleolin with iSN04 inhibited proliferation through the genome-wide regulation of cell cycle-related gene expression in myoblasts [Shinji 2021]. Since cell cycle arrest is a mandatory process for promoting myogenic differentiation [Singh 2013, Wu 2024], iSN04 likely triggered the myogenic gene expression program also in ARMS cells that retain myogenic potential. Since nucleolin is overexpressed in various cancers and sarcomas, including RMS, anti-nucleolin treatment represents a promising therapeutic strategy for tumors [Prakash 2025, Willmer 2021]. Another anti-nucleolin aptamer, AS1411, has been evaluated for relapsed acute myeloid leukemia and renal cell carcinoma in clinical trials, exhibiting efficacy with mild or moderate side effects [Van den Avont 2023]. Experimentally, AS1411 was reported to suppress the growth of an ERMS cell line, RD [Willmer 2021]. Interestingly, the anti-ERMS effects of iSN04 and AS1411 vary among ERMS cell lines [Nohira 2022]. Although AS1411-induced myogenic differentiation has been confirmed in myoblasts, it has not yet been demonstrated in RMS cells [Shinji 2021]. Thus, iSN04 provides an alternative antinucleolin treatment option for tumors, which is novel in terms of its ability to induce myogenic differentiation.

The iSN04-induced upregulation of myogenic genes, including myogenin, embryonic MHC, myomaker, and Ca_v_β^1^, was significant in RH30 cells but not in NRS-1 cells. Given that myotube-like NRS-1 cells expressed MHC even under control conditions, the basal expression levels of myogenic genes in NRS-1 cells were higher than those in RH30 cells. Therefore, the iSN04-dependent fold changes in myogenic induction appeared to be more prominent in RH30 cells. Regardless of these distinct myogenic properties between NRS-1 and RH30 cells, iSN04 consistently and significantly modulated the mRNA levels of ID1, ID3, and CASZ1 in both cell lines. ID proteins not only inhibit the upstream pathways of myogenesis during muscle development, but are also overexpressed in various cancers to activate oncogenic pathways [Lasorella 2014]. It is thus plausible that the iSN04dependent downregulation of ID1 and ID3 contributed to the induction of myogenic differentiation and the suppression of proliferation in NRS-1 and RH30 cells. To clarify how iSN04 regulates ID gene transcription, the involvement of nucleolin in this process requires further investigation. CASZ1 is a recently identified transcription factor that induces the differentiation of skeletal muscle and RMS cells. A previous study reported that the overexpression of CASZ1 in RH30 cells suppressed proliferation and induced myogenic differentiation by upregulating MyoD, myogenin, and MHC [Liu 2020 NC]. These findings correspond well with our observation that iSN04-induced CASZ1 expression and the subsequent upregulation of myogenic genes resulted in phenotypic changes in NRS1 and RH30 cells. The relationship between nucleolin and CASZ1 expression remains unknown. The mechanism underlying the iSN04-orchestrated gene expression shift in ARMS cells need to be uncovered through comprehensive analyses, such as transcriptomic profiling.

Combination experiments of iSN04 and a low dose of VCR, which quantified myogenic gene expression in RH30 cells, revealed that VCR has no effect in the absence of iSN04, and does not enhance the iSN04 activity even in the presence of iSN04. However, co-treatment with iSN04 and VCR cooperatively reduced the proliferation of RH30 cells compared with single-agent treatments. VCR targets tubulin dimers and inhibits microtubule formation, an anti-tumor mechanism of action that is independent of iSN04-induced myogenic differentiation. Therefore, the combined effect of iSN04 and VCR on the proliferation of RH30 cells is considered to be an additive rather than a synergistic effect. This suggests that iSN04 and VCR are unlikely to exacerbate their side effects. Combining iSN04 with VCR could also be beneficial in reducing the required dose of VCR, as highdose VCR treatment can increase the incidence of adverse events in RMS therapy [Madsen 2019, Carli 2004]. In clinical settings, VCR is administered in combination with ActD and cyclophosphamide (VCA) in North America, or with ActD and ifosfamide (IVA) in Europe [Miwa 2020]. ActD intercalates into GC-rich regions of double-stranded DNA and blocks RNA polymerases to inhibit transcription. Cyclophosphamide and ifosfamide form DNA-DNA cross-links and block DNA synthesis. The combined effects of iSN04 with these DNA-targeting agents on RMS cells should be evaluated in subsequent studies. Validating the efficacy and safety of iSN04 in combination with VCA and IVA using animal models would provide a novel therapeutic strategy integrating standard chemotherapy and differentiation therapy for RMS.

## 5. Conclusions

The anti-nucleolin aptamer iSN04 induces myogenic differentiation and suppresses the proliferation of two ARMS cell lines, NRS-1 and RH30. Co-treatment with iSN04 and VCR further inhibits the proliferation of RH30 cells compared with single-agent treatments. Thus, iSN04 holds potential for application in differentiation therapy for RMS, particularly in combination with current standard chemotherapy.

## Supporting information

iSN04 induces contracting myotube-like NRS-1 cells

## Supplementary Materials

Supplementary Video S1: iSN04 induces contracting myotube-like NRS-1 cells.

## Author Contributions

Conceptualization, T.T.; investigation, Y.S.; writing—original draft preparation, T.T.; funding acquisition, T.T. All authors have read and agreed to the published version of the manuscript.

## Funding

This research was funded by the Japan Society for the Promotion Science (JSPS) (25K09035) and the Japan Agency for Medical Research and Development (AMED) (JP24ym0126813j0003 and JP25ym0126819j0001) to T.T.

## Institutional Review Board Statement

Not applicable.

## Informed Consent Statement

Not applicable.

## Data Availability Statement

The data presented in this study are available on request to the corresponding author.

## Acknowledgments

The preprint has been posted on bioRxiv.

## Conflicts of Interest

The authors declare no conflict of interest.

## Abbreviations

ActD: Actinomycin D
ARMS: Alveolar rhabdomyosarcoma
Ca_v_β^1^: Voltage-gated calcium channel β1 subunit
ERMS: Embryonal rhabdomyosarcoma
ID: Inhibitor of DNA-binding / differentiation
IVA: Ifosfamide, vincristine, and actinomycin
DMEK: Mitogen-activated protein kinase kinase
MHC: Myosin heavy chain
MRF: Myogenic regulatory factor
RMS: Rhabdomyosarcoma
VCA: Vincristine, actinomycin
D,: and cyclophosphamide
VCR: Vincristine

## Notes

### Competing Interest Statement

The authors have declared no competing interest.

